# Introducing An Argonaute-facilitated Isothermal Amplification Technology

**DOI:** 10.1101/2025.04.21.649904

**Authors:** Feng Gao, Chun-Yu Han

## Abstract

Argonaute-facilitated targeting, which is an enzyme-mediated, high-fidelity, and highly efficient base pairing process, can be repurposed for upgrading PCR technology. An Argonaute protein derived from thermophilic *Caloramator sp*. (CalAgo) cleaved target dsDNA in the presence of Tte UvrD helicase at 65°C, suggesting dmCalAgo (a nuclease-inactive mutant) binding to target dsDNA in the same condition. Based on this, we designed an Argonaute-facilitated isothermal PCR platform (Isothermal Ago-PCR) through synergy of dmCalAgo, Tte UvrD helicase, and Bst DNA polymerase. Isothermal Ago-PCR realized amplification of template at 65°C. The upgrade resides in replacing primer annealing with Argonaute-facilitated targeting. Hence, Isothermal Ago-PCR not merely eliminates the reliance on complex primer design and sophisticated instruments but also achieves high-fidelity and highly efficient amplification. This platform enabled detection of low-copy templates (as few as 3 copies per reaction) within 30 minutes, achieving comparable sensitivity to qPCR while demonstrating superior amplification kinetics. This platform also demonstrated compatibility with common thermal maintenance tools. Thus, Isothermal Ago-PCR has the potential to replace qPCR with broad applicability. Details refer to our patent (China Patent CN116479095A).

## Results

An Argonaute protein was screened and designated as CalAgo from the thermophilic bacterium *Caloramator* sp. *ALD01* (NCBI Sequence ID: WP_027308646.1), by comparative sequence analysis of DNA-guided DNA pAgos (including TtAgo, NgAgo, and CbAgo)^[1, 2]^ (Figure 1A). The *in vitro* cleavage assays revealed that CalAgo possesses DNA-guided endonuclease activity whereas the dmCalAgo (a nuclease-inactive mutant) abolishes detectable endonuclease capability (Figure 1B, 1C)^[3, 4]^. Furthermore, we systematically characterized the enzymatic properties of CalAgo. In terms of divalent metal cation dependence, CalAgo exhibited cleavage activity in the presence of Mg^2+^, Mn^2+^, Co^2+^, or Ca^2+^, with optimal efficiency observed for Mg^2+^ and Mn^2+^ (Figure 1D). Temperature profiling revealed that CalAgo maintained cleavage activity across a broad thermal range (37–85°C), with the activity peaking between 55°C and 75°C (Figure 1E). As for gDNA length requirements, CalAgo demonstrated robust cleavage activity with 14nt-30nt gDNA and retained detectable cleavage activity even with 50nt gDNA (Figure 1F).

**Fig 1.**
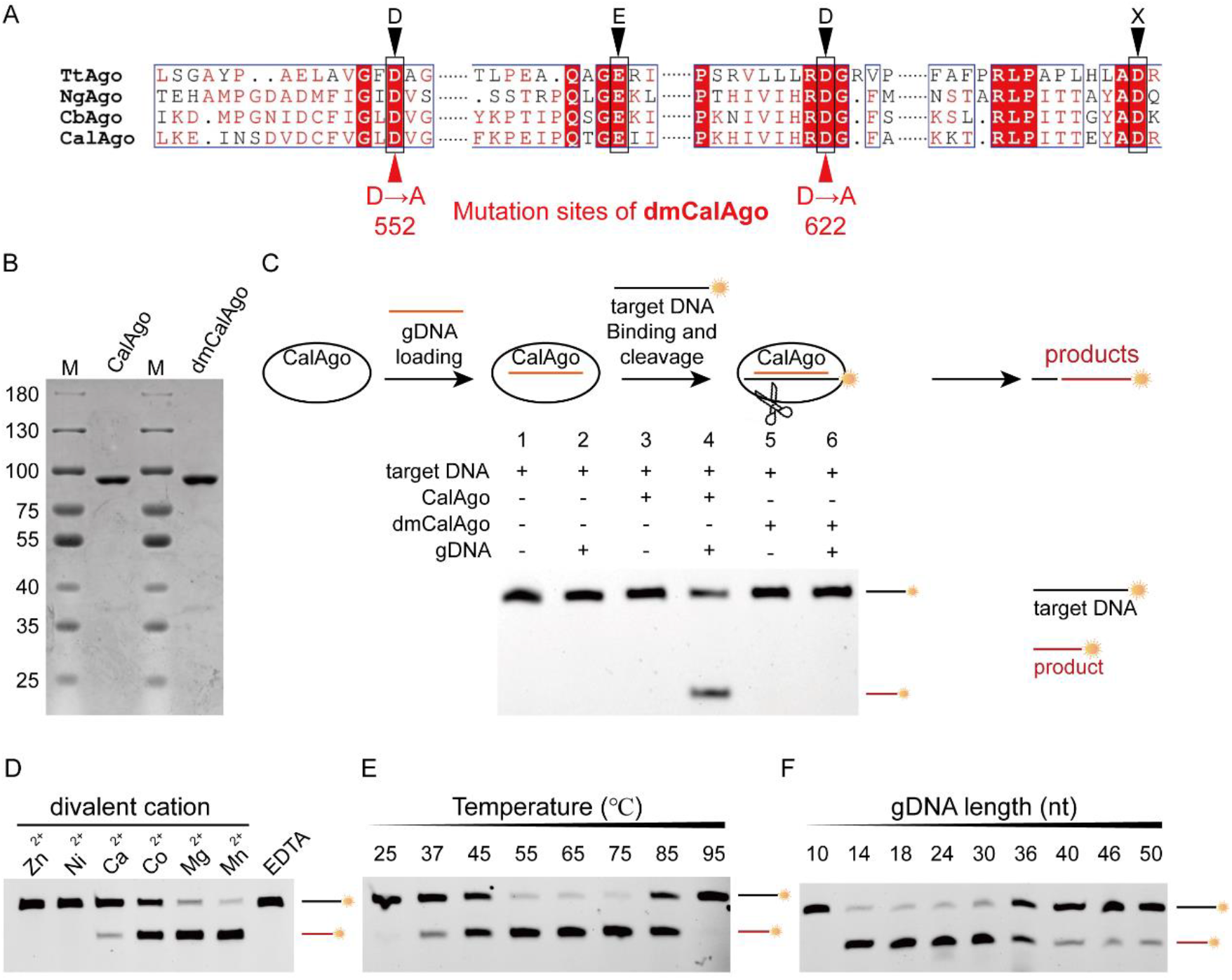
CalAgo exhibits DNA-guided DNA endonuclease activity. **(A)** Sequence alignment of conserved amino acid residues of the DEDX tetrad (marked by black triangles) within the PIWI domains of CalAgo, TtAgo, NgAgo, and CbAgo. A nuclease-inactive mutant of CalAgo (designated as dmCalAgo) was engineered with double mutated sites (D552A and D622A, marked by red triangles). **(B)** SDS-PAGE of purified CalAgo and dmCalAgo (Ni-NTA affinity chromatography). **(C) Upper:** Scheme of the *in vitro* assay. All target DNA substrates were labeled with Alexa Fluor 488 at the 5′-end (marked by yellow sun). **Lower:** CalAgo exhibites DNA-guided ssDNA endonuclease activity while dmCalAgo loses the endonuclease activity. The gDNA used in these experiments was pLGC-18. Sequences of Target DNA and gDNA are detailed in Materials and Methods. **(D)** The contribution of various divalent cations on DNA-guided ssDNA endonuclease activity of CalAgo. **(E)** The effect of temperature on ssDNA cleavage by CalAgo. The target ssDNA was added after the CalAgo and gDNA had been incubated for 15 minutes at 65°C. Then the integral reaction mixture was incubated for 30 minutes at various temperatures. **(F)** The dependence of gDNA length on DNA cleavage activity by CalAgo. The guides used in these experiments were pLGC-10, pLGC-14, pLGC-18, pLGC-24, pLGC-30, pLGC-36, pLGC-40, pLGC-46, pLGC-50. Sequences of guides are detailed in Materials and Methods.

In the presence of Tte UvrD helicase, which is thermostable and unwinds dsDNA efficiently at 65°C, CalAgo exhibited robust dsDNA cleavage activity (Figure 2). This synergistic effect likely arises from the helicase-mediated unwinding of dsDNA into transient ssDNA regions, which subsequently allows sequence-specific binding and cleavage by the CalAgo/gDNA complex.

**Fig 2.**
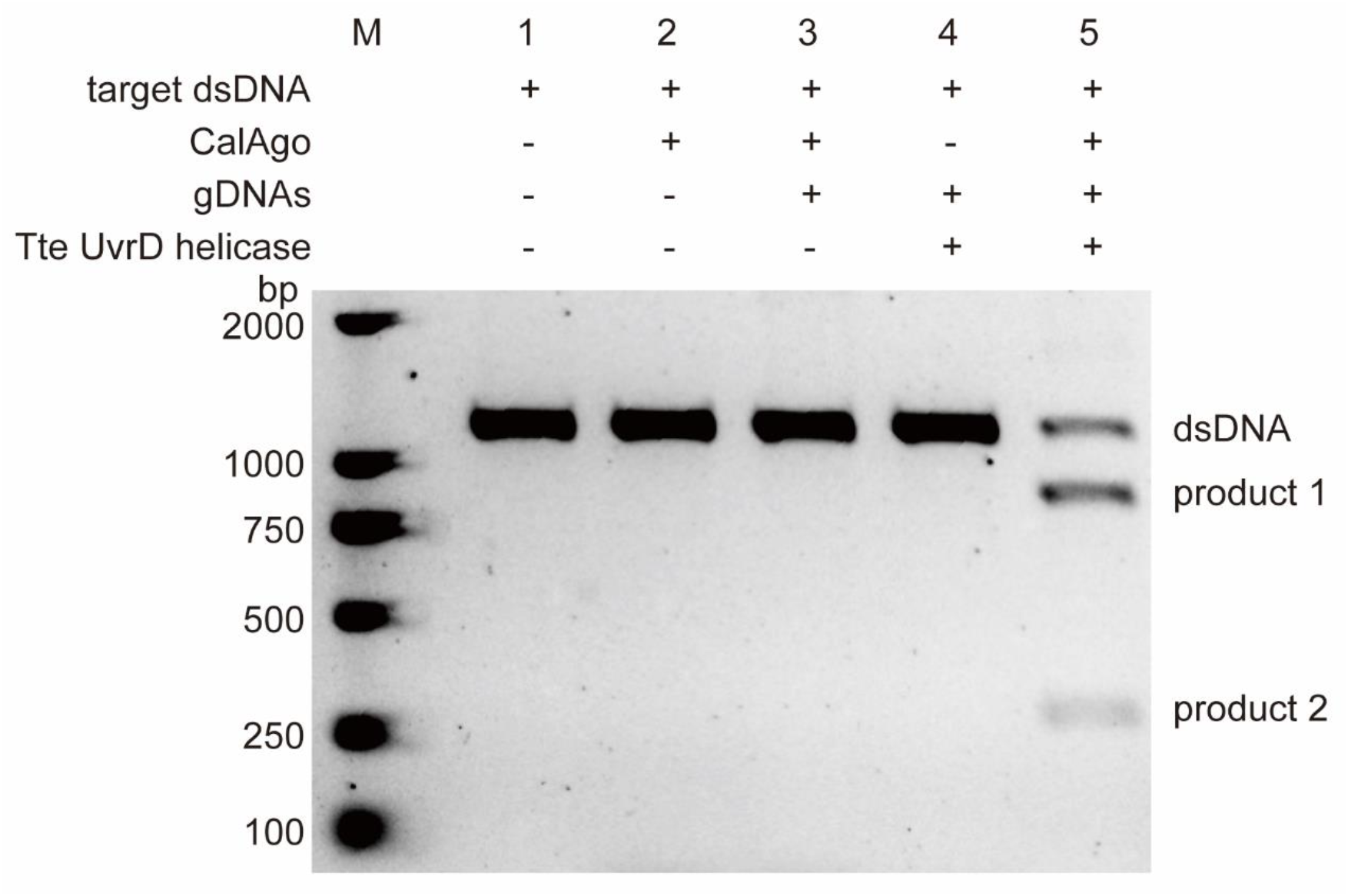
CalAgo cleaves target dsDNA in the presence of Tte UvrD helicase at 65°C. The guides used in these experiments were pLGC-18 and T3-pLGC-18.

Therefore, we reasoned that this synergistic effect could be repurposed for upgrading annealing in isothermal amplification^[5, 6]^. By combining the cleavage-deficient dmCalAgo mutant and Tte UvrD helicase with a strand-displacing DNA polymerase (e.g., Bst LF)^[7]^, we propose an Argonaute-facilitated isothermal amplification platform, termed Isothermal Ago-PCR (Figure 3).

**Fig 3.**
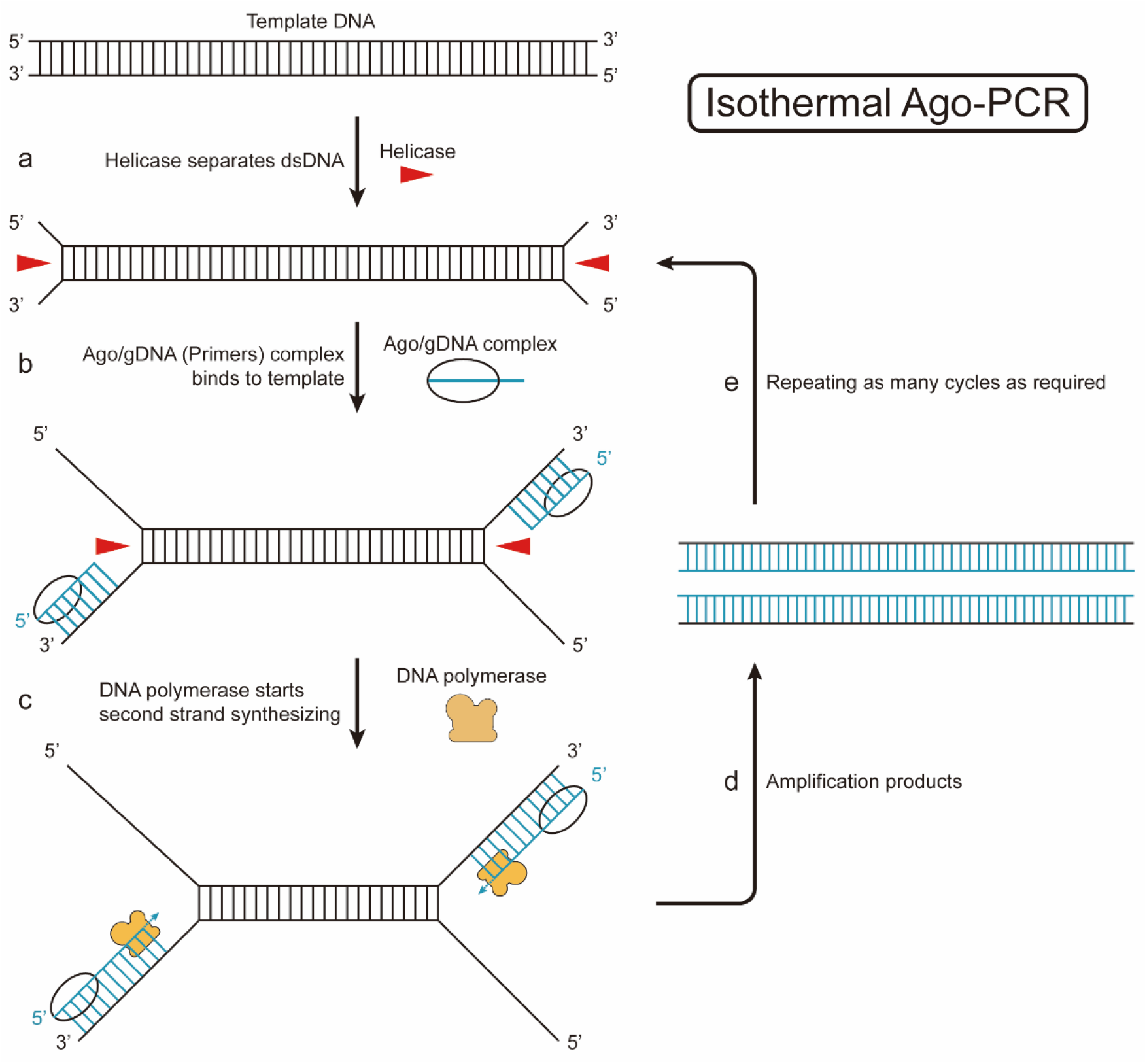
The theoretical working model for Isothermal Ago-PCR. In this model: **a**. Tte UvrD helicase helps to unwind the target dsDNA into transient ssDNA. **b**. dmCalAgo protein/guide DNA complexes target the target DNA and execute sequence-specific binding, with the 3 ‘end of gDNA (functioning as primer) exposed. **c**. DNA polymerase recognizes the exposed 3’ end of gDNA and initiates polymerization, while target DNA are used as templates. **d**. Generating amplification products which act as the templates for the next cycle. **e**. Cycling through a-d steps enables exponential amplification of target templates.

Isothermal Ago-PCR platform demonstrated robust amplification efficiency for M13mp18 template, as evidenced by clear product accumulation in gel electrophoresis analysis (Figure 4).

**Fig 4.**
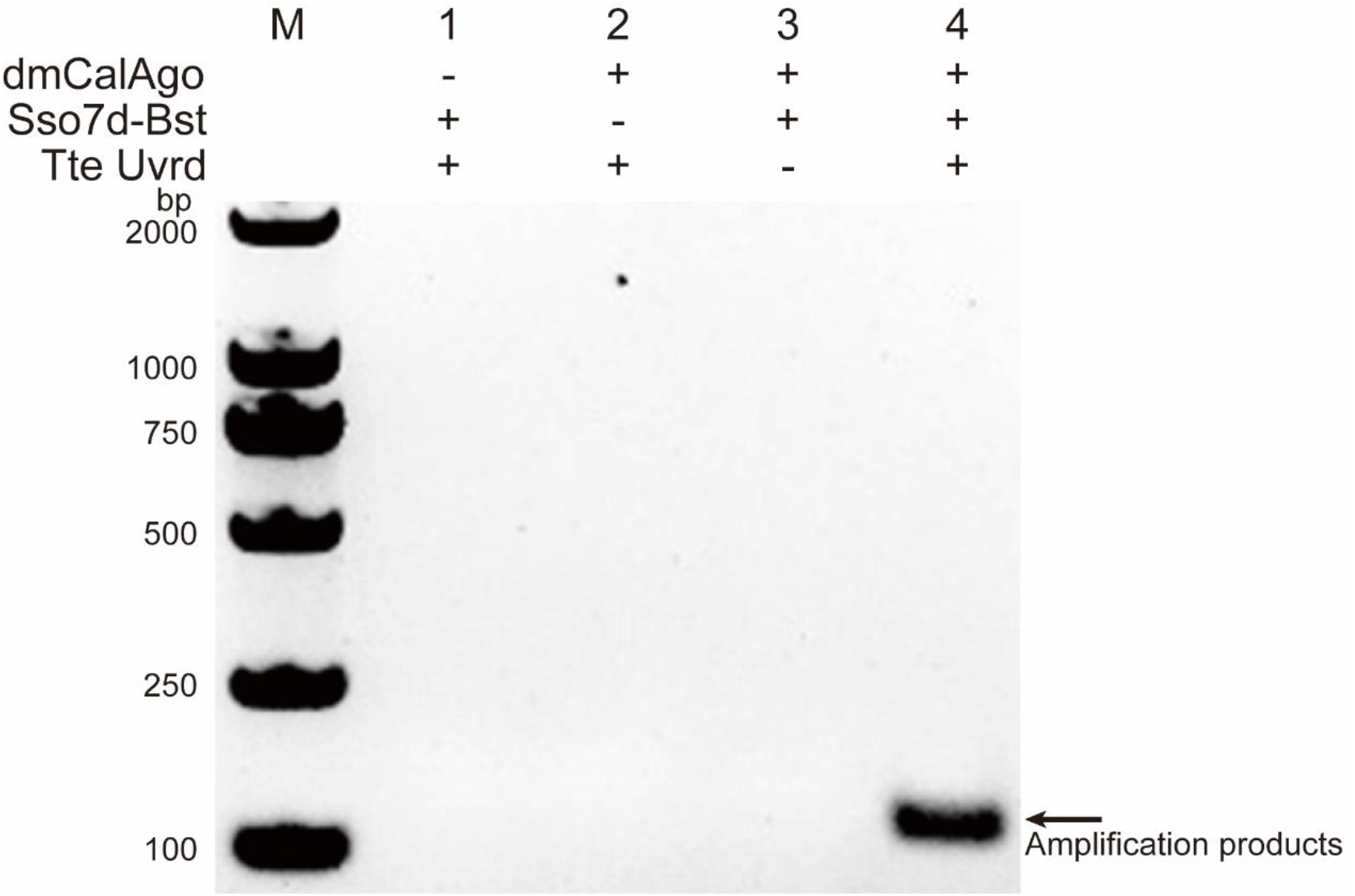
Demonstration of the isothermal Ago-PCR at 65°C. 6×10^4^ copies per reaction of M13mp18 template was used in these experiments. The guides (primers) used in these experiments were pM13-3-30 and pM13-7-40. The complete reaction mixture was incubated at 65°C for 30min. The products were resolved by 2.5% agarose gel.

Moreover, Isothermal Ago-PCR also achieved comparable amplification efficiencies observed across circular ssDNA, linear dsDNA, and plasmid, exhibiting broad template versatility (Figure 5). These results strongly support the feasibility of our mechanistic model.

**Fig 5.**
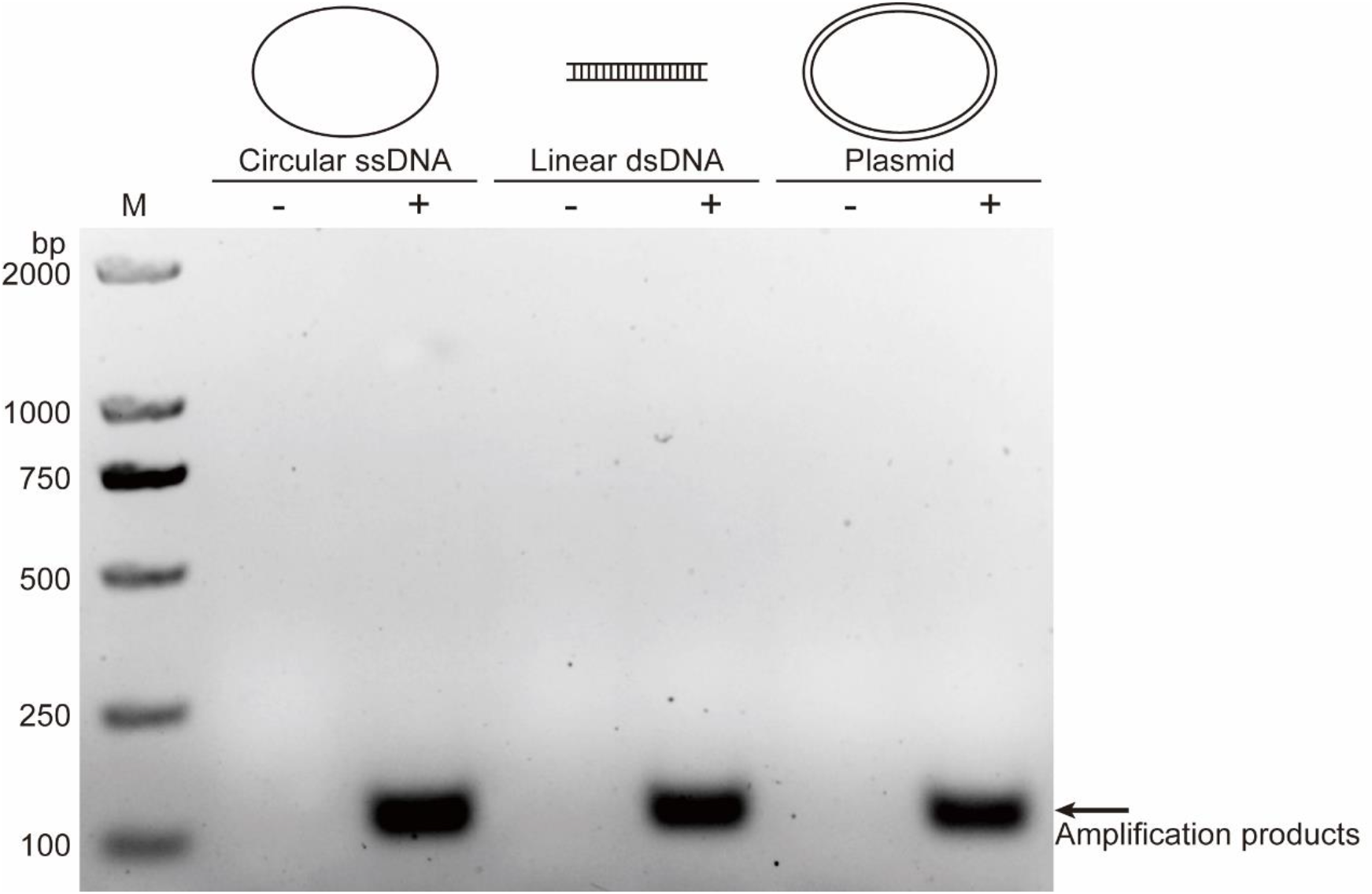
Isothermal Ago-PCR successfully amplifies various DNA substrates. Circular single-strand DNA M13mp18 template (circular ssDNA), M13mp18 vector-based linear dsDNA template (linear dsDNA), and circular double-stranded DNA pM13 template (plasmid) were successfully amplified in Isothermal Ago-PCR. 300 copies of each template per reaction were used in these experiments. The guides (primers) used in these experiments were pM13-3-30 and pM13-7-40.

A benchmarking analysis comparing the sensitivity and amplification kinetics between Isothermal Ago-PCR and conventional qPCR revealed striking performance advantages of the former. Quantitative assessments demonstrated that Isothermal Ago-PCR achieves equivalent sensitivity (3 copies per reaction) to qPCR with significantly accelerated reaction kinetics (Figure 6).

**Fig 6.**
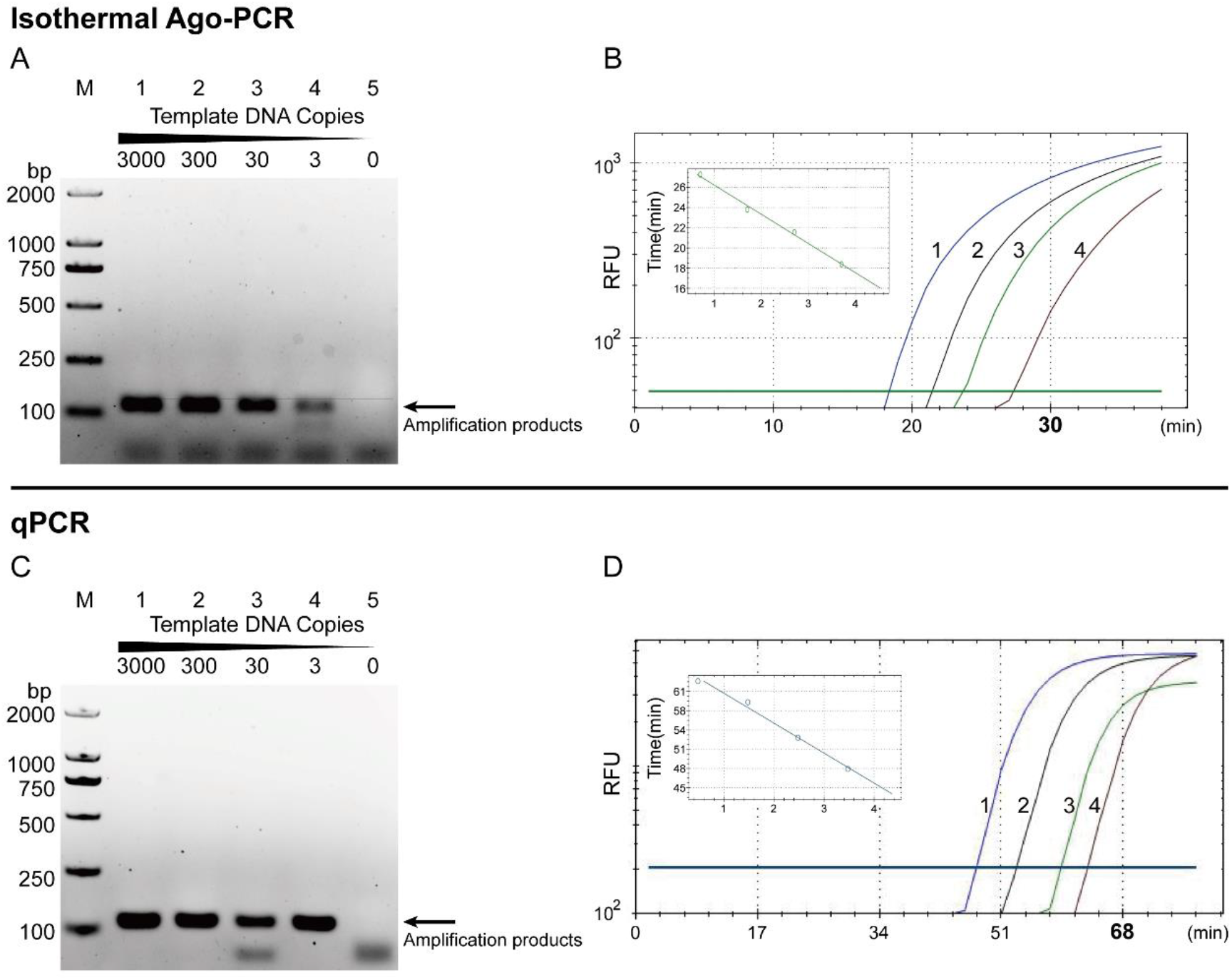
Comparison of sensitivity and amplification efficiency between isothermal Ago-PCR and qPCR. (A-B) Isothermal Ago-PCR detected various copies of M13mp18. The results were examined by (A) argrose electrophoresis, and (B) real-time fluorescence detection (SYBR Green I). Isothermal Ago-PCR achieved detection of 3 copies per reaction of M13mp18 template within 30 minutes. (C-D) qPCR detected various copies of M13mp18. The results were examined by (C) argrose electrophoresis, and (D) real-time fluorescence detection (Taqman probe). qPCR achieved detection of 3 copies per reaction of M13mp18 template beyond 60 minutes.

Critically, Isothermal Ago-PCR achieved sensitive detection of low-copy targets (<10 copies per reaction) using only common thermal maintenance tools (Figure 7). These streamLined hardware requirements position the platform as a field-deployable solution for point-of-care diagnostics.

**Fig 7.**
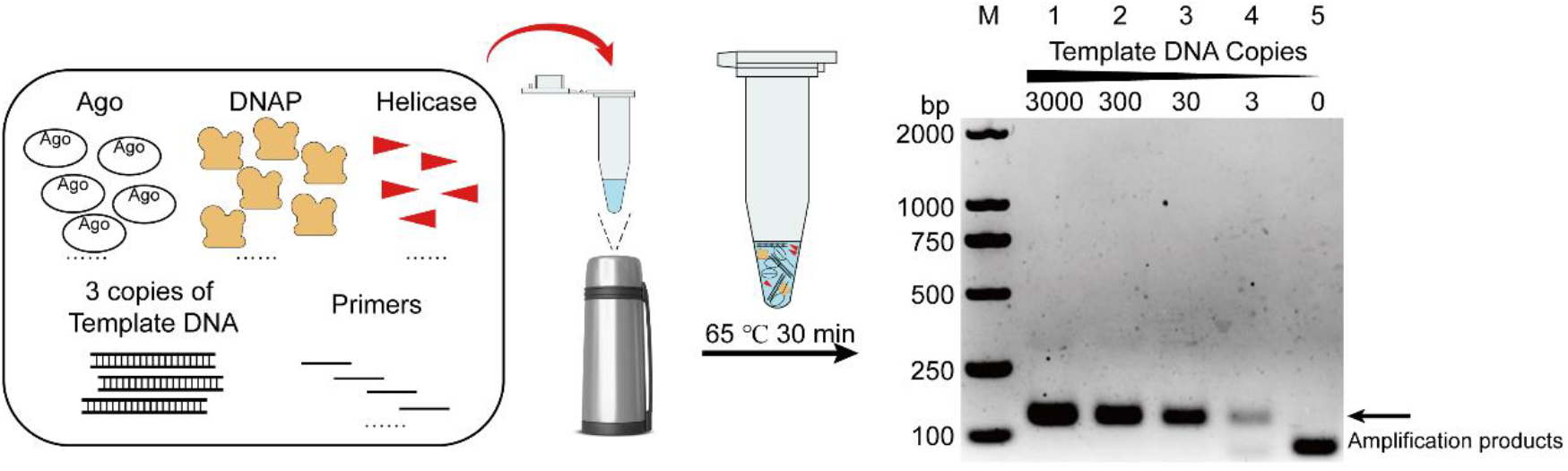
Isothermal Ago-PCR successfully detects 3 copies of templates without the need for sophisticated equipment.

## Acknowledgements

We thank Xiao Z. Shen (Department of Physiology, Zhejiang University School of Medicine, Hangzhou, Zhejiang, China.), Feng Jiang, Jianhui Yang, Ruoqing Liu, Yinuo Zhang, Zizheng Li, Xingyu Dong, Peijiao Sun for the contributions.

## Materials and Methods

### Protein expression and purification

The CalAgo gene (*Caloramator sp. ALD01*, Sequence ID: WP_027308646.1), double mutant CalAgo (dmCalAgo) (D552A and D622A) gene, sso7d-Bst DNA polymerase gene (*Bacillus stearothermophilus*, GenBank:ALA69666.1,sso7d protein fused to the N-terminal of Bst DNA polymerase large fragment) and Tte UvrD gene (*Thermoanaerobacter tengcongensis MB4*, GenBank: AAM23874.1) were synthesized by Shanghai Sangon Biotech and cloned into pET-28a(+) expression vectors in frame with the N-terminal 6×His tag. The recombinant proteins expression and Ni-affinity chromatography purification procedures were performed according to user manual (Novagen). In particular, the resultant purified proteins were eluted with elution buffer (20 mmol/L Tris-HCl pH 7.5, 500 mmol/L NaCl, 200 mmol/L Imidazole), and concentrated with 50Kda column (Merck Millipore#UFC905008) to 1μg/μL. Adding glycerol to final concentration of 50% for preservation.

### DNA guided Target DNA cleavage assays

Unless stated otherwise CalAgo, gDNA and Target DNA were mixed in a molar ratio of 5:2:1, in reaction buffer (20mM Tris-HCl, 50mM KCl, 10mM MgCl_2_, pH 7.5). The target DNA was added after the CalAgo and gDNA had been incubated for 15 min at 65°C. Then the complete reaction mixture was incubated for 30min at 65°C. The reaction was stopped by 2×RNA loading buffer (1×TBE, 6% Ficoll^®^-400, 5mM EDTA, 0.05% SDS,85% Formamide) and heating it for 5min at 95°C. The cleavage products were resolved by 15% denaturing polyacrylamide gel. The gel was imaged using ChemiDoc™ Touch Imaging System (BIO-RAD). Sequences of Target DNA and gDNA are detailed in Table.

### CalAgo-dsDNA cleavage assays

500nM CalAgo protein, 250nM gDNA (pLGC-18) and 250nM gDNA (T3-pLGC-18) were preincubated at 65°C in reaction buffer (20mM Tris-HCl, 50mM KCl, 10mM MgCl_2_, pH 7.5) for 15min. Then add 50nM Target dsDNA, 1μg Tte UvrD, 2.5nM dATP, add water to final volume to 20μL. The complete reaction mixture was incubated at 65°C for 30min. The reaction was stopped by 1mg/mL Proteinase K for 1h at 60°C. The cleavage products were resolved by 1% agarose gel.

The gel was imaged using ChemiDoc™ Touch Imaging System (BIO-RAD). Target DNA was synthesized by Shanghai Sangon Biotech and cloned into pGEM^®^-T Easy (PROMEGA #A137A). The resultant plasmid is designated as pTarget-DNA. Target dsDNA was produced by amplifying pTarget-DNA by using primers pTarget-F-55 and pTarget-R-56. The resultant PCR product is 1210bp.

### Isothermal Ago-PCR

300ng CalAgo,60nM pM13-3-30 and 60nM pM13-7-40 were preincubated at 65°C in reaction buffer (20mM Tris-HCl, 10mM (NH_4_)_2_SO_4_, 150mM KCl, 2mM MgSO_4_, 0.1% Tween^®^20, 1mM DTT, pH 8.8) for 5min, then add DNA template, 160μM dNTP Mixture, 100ng sso7d-Bst DNA polymerase, 200ng Tte UvrD, 2nM dATP add water to final volume to 30μL. The complete reaction mixture was incubated at 65°C for 30min. The products were resolved by 2.5% agarose gel. The gel was imaged using ChemiDoc™ Touch Imaging System (BIO-RAD).

### qPCR

Reaction system: 200nM forward primer (M13-4-F-22), 200nM reverse primer (M13-4-R-20), probe (F-M13-2), 200μM dNTP, 1U Taq DNA Polymerase (NEB), 1X Standard Taq Reaction Buffer (NEB B9014S) and certain dosage of M13mp18 template according needs, add water to final volume to 20μL. Program: 95°C 3min, 95°C 15s, 59°C 30s, 68°C 10s plate read, step 5 Go to step 2, 45 cycles, 68°C 5min end.

### Template preparation

M13mp18 template (NEB #N4040S) is the 7249nt circular single-stranded DNA and obtained from New England Biolabs. The 4330bp PCR product amplified from M13mp18 by using primers: oM13-5-27 and oM13-3-24 was inserted into pGEM^®^-T Easy (PROMEGA #A137A). The resultant plasmid is designated as pM13. pM13 is used as double stranded, circular template. The double stranded, linear template (dsDNA4330) was produced by amplifying M13mp18 by using primers oM13-5-27 and oM13-3-24. The resultant PCR product is 4330bp.

**Table 1.**
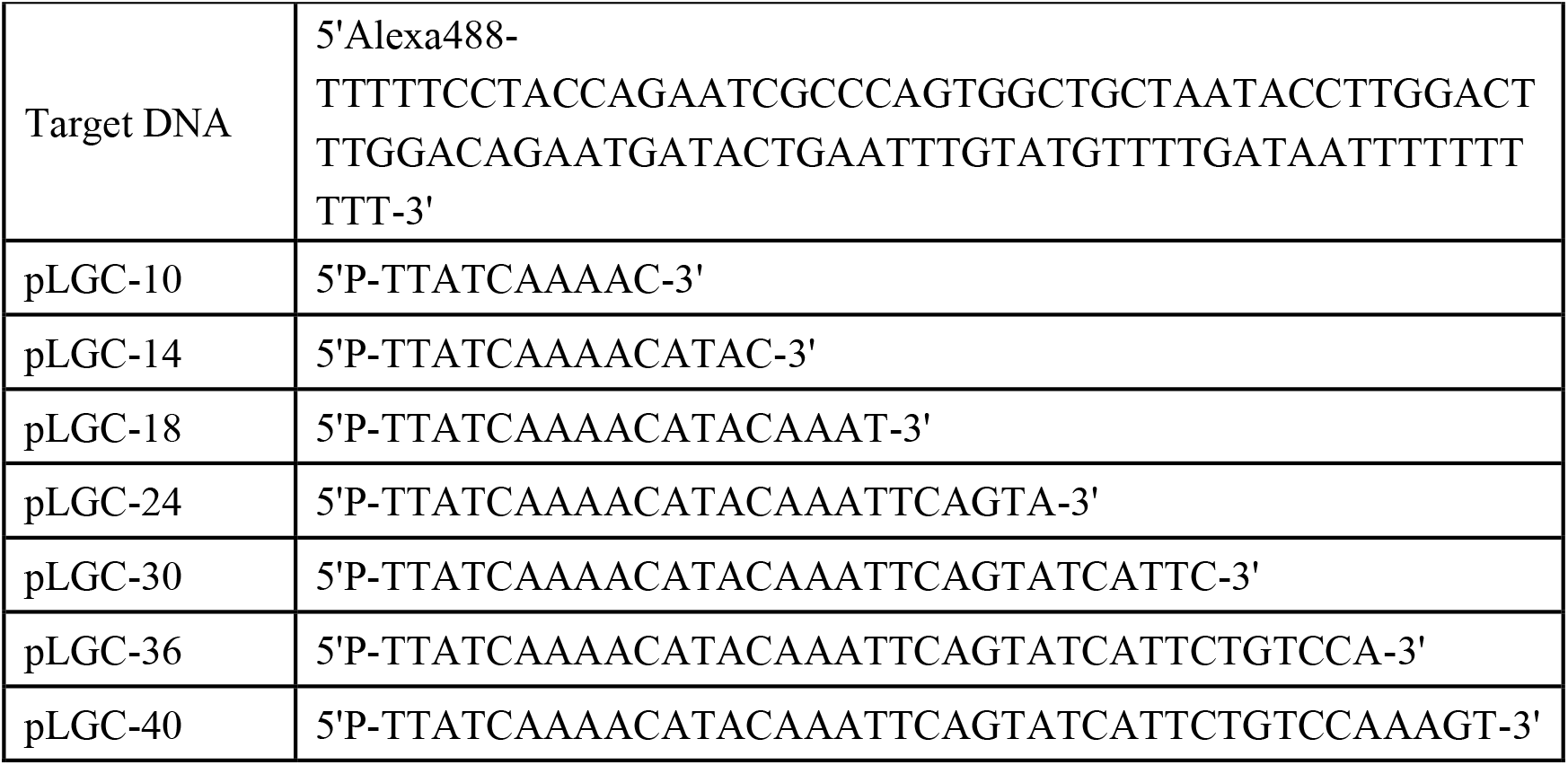
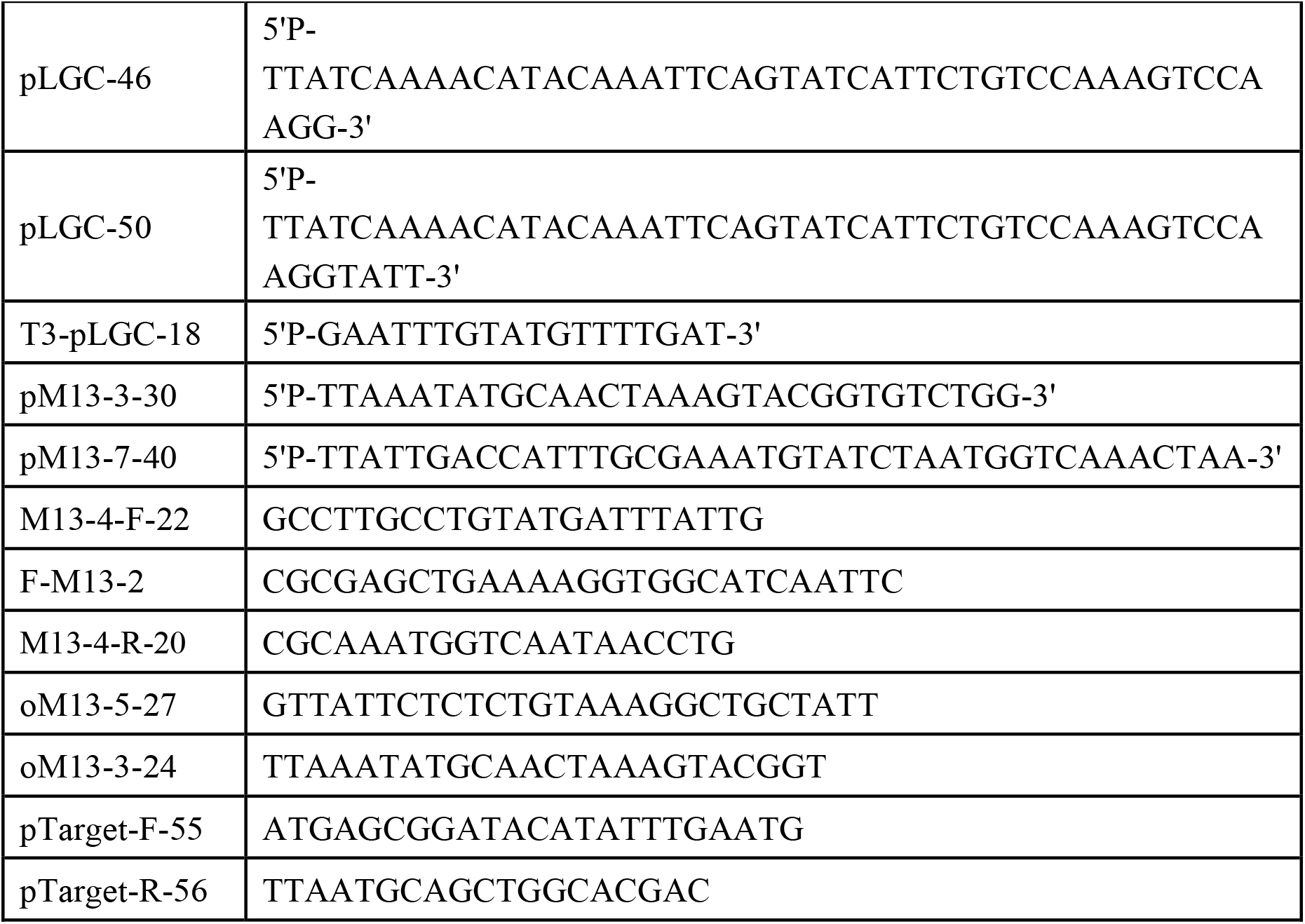
Sequences of guides and primers

